# Diversity within olfactory sensory derivatives revealed by the contribution of Dbx1 lineages

**DOI:** 10.1101/2022.11.12.514524

**Authors:** Frédéric Causeret, Maxime Fayon, Matthieu X. Moreau, Enrico Ne, Roberto Oleari, Carlos Parras, Anna Cariboni, Alessandra Pierani

## Abstract

In vertebrates, the embryonic olfactory epithelium contains progenitors that will give rise to distinct classes of neurons, including olfactory sensory neurons (OSN, involved in odor detection), vomeronasal sensory neurons (VSN, responsible for pheromone sensing) and GnRH neurons that control the hypothalamic-pituitary-gonadal axis. Currently, these three neuronal lineages are usually believed to emerge from uniform pools of progenitors. Here we found that the homeodomain transcription factor Dbx1 is expressed by neurogenic progenitors in the developing and adult mouse olfactory epithelium. We demonstrate that Dbx1 itself is dispensable for neuronal fate specification and global organization of the olfactory sensory system. Using lineage tracing we characterize the contribution of Dbx1 lineages to OSN, VSN and GnRH neuron populations and reveal an unexpected degree of diversity. Furthermore, we demonstrate that *Dbx1*-expressing progenitors remain neurogenic in the absence of the proneural gene *Ascl1*. Our work therefore points to the existence of distinct neurogenic programs in Dbx1-derived and other olfactory lineages.

## Introduction

Vertebrates share an olfactory sensory system specialized in odor detection and pheromone sensing (Poncelet and Shimeld, 2020). Components of the olfactory sensory system emerge from the olfactory placode (OP) that progressively thickens and differentiate into a morphologically distinct olfactory epithelium (OE) from early steps of embryonic development (around E9.5 in mice). Olfactory sensory neurons (OSN) are the main output of the OE, they extend multiple cilia bearing transmembrane odorant receptors in the nasal cavity, and project their axon to the main olfactory bulb (OB). Vomeronasal sensory neurons (VSN), involved in pheromone sensing, arise from a specialized part of the OE and project to the accessory OB (Katreddi and Forni, 2021). Neurogenesis of OSN and VSN is a continuous process, starting around E10.5 in mice and extending throughout adulthood, therefore allowing constant cell renewal (Sokpor et al., 2018). The distinct cellular states through which OSN and VSN transit during differentiation are relatively well characterized, and include self-renewing progenitors, intermediate progenitors committed to neuronal fate, immature neurons and eventually mature OSN/VSN characterized by the expression of the olfactory marker protein (OMP) (McClintock et al., 2020; Schwob et al., 2017). In adults, the differentiation trajectories leading to the production of OSN and lineage relationship with glial components of the OE have been precisely resolved through scRNAseq (Fletcher et al., 2017; Hanchate et al., 2015).

In addition to OSN and VSN, the embryonic OP also gives rise to gonadotropin-releasing hormone (GnRH) neurons (Duan and Allard, 2020; Wray et al., 1989). Upon specification at the vicinity of the vomeronasal organ (VNO) anlage around E11.5 in mice, GnRH neurons delaminate from the OP and engage a long range migration in the nasal mesenchyme towards the basal forebrain, to eventually settle in the hypothalamus (Cho et al., 2019). These neurons are of critical physiological importance, as GnRH secretion regulates the release of luteinizing and follicle-stimulating hormones from the pituitary gland, and subsequent production of sexual steroids from the gonads thus regulating sexual maturation and reproductive function. Consequently, defects in the specification or migration of GnRH neurons result in hypogonadotropic hypogonadism, as observed in Kallmann syndrome (Oleari et al., 2021). Some OP derivatives remain poorly characterized. This is the case of the so-called “migratory mass” (MM), a heterogeneous population including neurons and glial cells, strategically positioned in between the OE and ventral telencephalon. To date, the precise composition, origin and function of the MM is still poorly defined, although a role as guidepost cells for pioneer OSN axons was proposed (Miller et al., 2010; Valverde et al., 1992).

The molecular logic of neuronal specification in the olfactory sensory system is being increasingly understood and appear similar at embryonic and adult stages (Sokpor et al., 2018). First, progenitors pools express and require the transcription factor Sox2 for their maintenance and proliferation (Donner et al., 2007). Commitment to neuronal differentiation involves the transition to intermediate progenitor state and is regulated by the successive expression of *Ascl1* and *Neurog1* (Cau et al., 1997; Cau et al., 2002; Guillemot et al., 1993). Finally, lineage-dependent maturation programs unfold and enable appropriate neuronal maturation. Although the main transcription factors expressed in the OE/VNO and involved in fate specification have been described (Chang and Parrilla, 2016; Parrilla et al., 2016), some remain uncharacterized. This is the case of Dbx1 which is the main focus of this study. Dbx1 is a homeodomain transcription factor whose expression is largely restricted to the developing nervous system (Lu et al., 1992; Shoji et al., 1996). In the spinal cord, hindbrain and hypothalamus, *Dbx1* is involved in neuronal fate specification (Bouvier et al., 2010; Pierani et al., 2001; Sokolowski et al., 2015). We show that *Dbx1* transcript and protein are dynamically expressed during embryonic and adult olfactory neurogenesis. Reminiscent of the central nervous system, *Dbx1* expression is spatially and temporally constrained to committed neural progenitors. We found *Dbx1* expression to be dispensable for neuronal production and global organization of the olfactory sensory system. We performed lineage tracing to follow the progeny of Dbx1-expressing progenitors and surprisingly found a contribution to OSN, VSN, GnRH and MM neurons, therefore revealing an unanticipated degree of diversity within these populations. Finally, we examined *Ascl1* mutants to reveal that unlike all other known olfactory progenitors, *Dbx1*-epxressing cells remain neurogenic in these animals. Our work therefore points to previously unknown mechanism of neuronal specification and diversification in the developing olfactory sensory system.

## Material & Methods

### Animals

All mouse work was approved by the French Ministry of Higher Education, Research and Innovation as well as the Animal Experimentation Ethical Committee of Paris Descartes University (CEEA-34, licence number: 18011-2018012612027541). The following mouse lines were used: *Dbx1*^*LacZ*^ (Pierani et al., 2001), *Dbx1*^*Cre*^ (Bielle et al., 2005), *Rosa26*^*LoxP-Stop-LoxP-YFP*^ (Srinivas et al., 2001), *Tau*^*GFP-NLS-LacZ*^ (Hippenmeyer et al., 2005) and *Ascl1 (Mash1)* knock-out (Guillemot et al., 1993).

### Immunostaining and *in situ* hybridization

Samples were fixed by immersion in 4% paraformaldehyde in 120mM phosphate buffer (PB) for 1-4h. They were cryoprotected overnight in 10% sucrose in PB and embedded in 7.5% Gelatin, 10% sucrose in PB. 20μ-thick cryostat sections were performed and processed for *in situ* hybridization, immunostaining or X-Gal staining (Bielle et al., 2005; Causeret et al., 2011). *In situ* hybridization probes for *Dbx1, Ascl1* (*Mash1), Neurog1* (*Ngn1)*, and *Lhx2* were generated by *in vitro* transcription of linearized plasmids described elsewhere (Cau et al., 1997; Guillemot and Joyner, 1993; Lu et al., 1992; Rétaux et al., 1999). Probes for *Stmn2* (*Scg10*), *Gap43, Ncam1, Omp, Gnrh1, Vmn1r180* (*V1rd16*) and *Vmn2r121* were obtained from PCR fragments amplified using primers indicated in Supplementary Table 1. Because of sequence homology, both vomeronasal receptor probes are predicted to hybridize with multiple genes of their respective family and were therefore referred to as *Vmn1r* and *Vmn2r* probes in this study. Probes for olfactory receptors were obtained from PCR fragments amplified using previously published primers (Miyamichi et al., 2005).

Antibodies were used at the following concentrations: chick anti-GFP (Aves Labs GFP-1020, 1:2000), chick anti-β Galactosidase (Abcam ab9361, 1:1000), rabbit anti-Phospho Histone 3 (Cell Signaling Technology #3377, 1:500), rabbit anti-Cleaved Caspase-3 (Cell Signaling Technology #9664, 1:1000), rabbit anti-Dbx1 (Pierani et al., 1999), rabbit anti-GnRH (Immunostar #20075, 1:400), goat anti-Sox2 (R&D Systems AF2018, 1:100), goat anti-OMP (Wako #019-22291, 1:1000) rat anti-CTIP2 (Abcam ab18456, 1:300) mouse anti-Tubulin β3 (Tuj1, BioLegend #801201, 1:1000) rabbit anti-Peripherin (Sigma-Aldrich AB1530, 1:500), goat anti-Lhx2 (Santa-Cruz sc-19344), mouse anti-HuC/D (Clone 16A11, ThermoFisher A-21271). Secondary antibodies coupled to Alexa488, Cy3, Cy5 or HRP were purchased from Jackson ImmunoResearch.

### Imaging

Visible staining was acquired either on a Zeiss Axiovert microscope or a Keyence VHX-2000 microscope. Fluorescent images were obtained with Zeiss LSM710 or Leica SP5 confocal microscopes.

## Results

### Dbx1 is expressed in the olfactory epithelium

*Dbx1* was previously reported to be expressed in the olfactory placode at E9.5 in mouse (Causeret et al., 2011). Given its well-characterized function in neuronal fate specification (Bouvier et al., 2010; Pierani et al., 2001; Sokolowski et al., 2015), we thought to evaluate its putative involvement in olfactory neurogenesis. We performed *in situ* hybridization experiments on coronal cryosections of mouse embryos at E10.5 and E11.5, stages corresponding to the onset of neuronal production. We found *Dbx1* to be expressed by scattered cells that preferentially occupied medial or dorso-lateral positions within the olfactory epithelium (OE) at E10.5 (Fig 1A). A similar distribution was found at E11.5 with most *dbx1*-expressing cells being found in the anterior and medial aspect of the OE and only few in dorso-lateral regions (Fig 1A). At this stage, *Dbx1* expression was not detected in the caudal half of the OE (not shown). Immunostaining confirmed that the Dbx1 protein is expressed in the OE (Fig 1B). We then used the *Dbx1*^*LacZ/+*^ mice in which expression of β-galactosidase (βGal) was previously shown to faithfully recapitulate that of *Dbx1* (Pierani et al., 2001). X-Gal staining indicated that Dbx1 expression decreased from E12.5 to E14.5, with only few cells found within the developing OE, and increased again at E18.5 (Fig 1C). Together, these data indicate that *Dbx1* is dynamically expressed in the embryonic olfactory epithelium.

**Figure 1.**
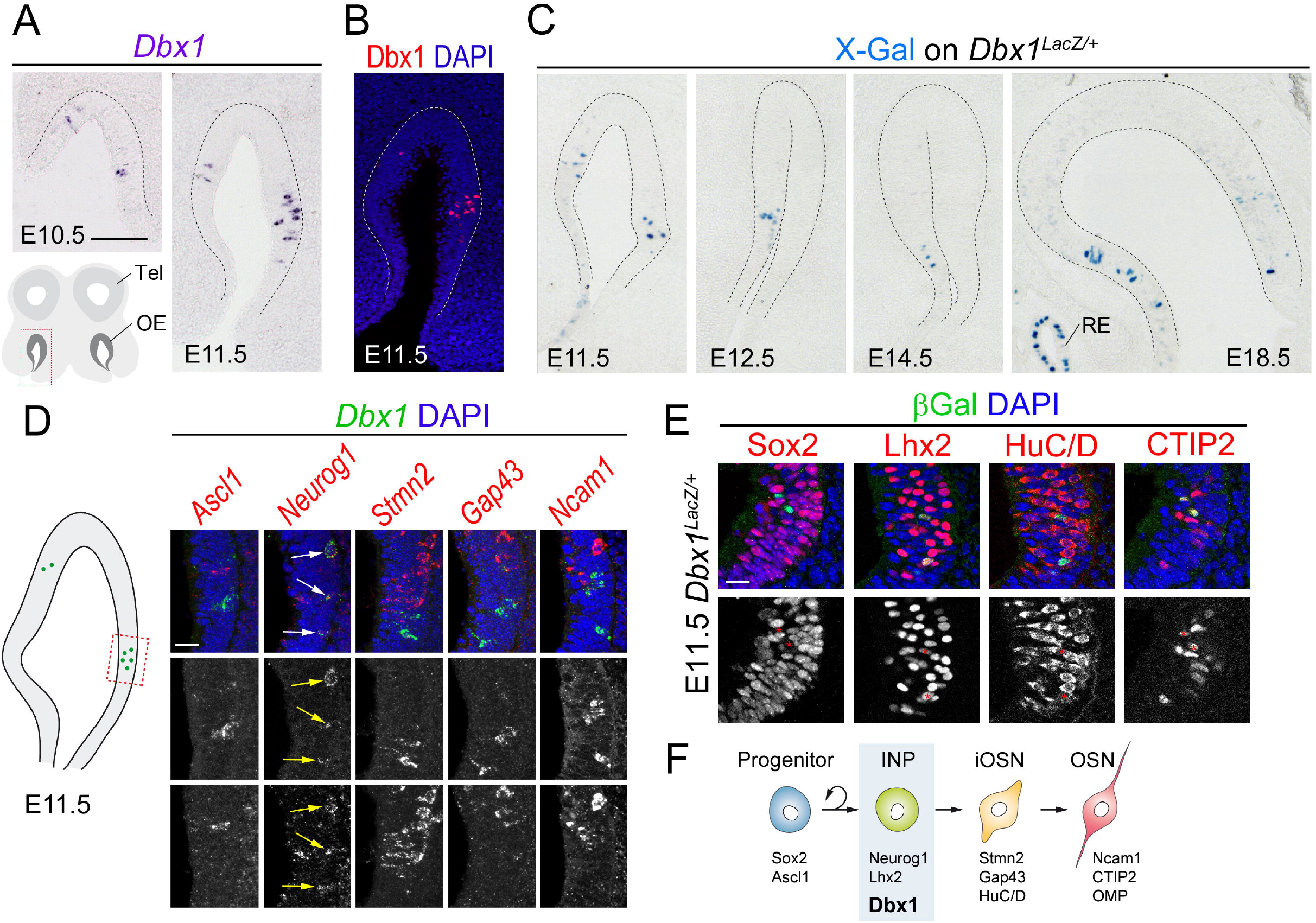
Expression of Dbx1 during olfactory development. (A) *In situ* hybridization using a probe against Dbx1 on coronal sections of the E10.5 and E11.5 mouse olfactory pit. The bottom left drawing indicates the level at which sections were collected. (B) Immunostaining for Dbx1 (red) on sections similar to (A). Nuclei are stained using DAPI (blue) (C) X-Gal staining on coronal sections from E11.5 to E18.5 *Dbx1*^*LacZ/+*^ embryos. (D) High magnification of a region of the E11.5 olfactory epithelium containing Dbx1-expressing cells (as indicated on the drawing) following double fluorescent *in situ* hybridization with probes against *Dbx1* (green) and the indicated genes (red). Nuclei are shown in blue using DAPI. (E) Immunofluorescent staining on E11.5 *Dbx1*^*LacZ/+*^ embryos using antibodies against βGalactosidase (green) and the indicated proteins (red) and DAPI (blue) to stain nuclei. (F) Drawing summarizing the findings: Dbx1 is expressed by neurogenic progenitors. OE: olfactory epithelium, Tel: telencephalon, RE: respiratory epithelium. INP: intermediate neural progenitor, OSN: olfactory sensory neuron, iOSN: immature OSN. Scale bars: 100μm in A, 20μm in D, E.

In the mouse developing brain and spinal cord, Dbx1 is expressed by neurogenic progenitors and subsequently downregulated in postmitotic neurons (Bielle et al., 2005; Lu et al., 1992; Pierani et al., 2001). In order to test whether this also applies to the olfactory system, we analyzed *Dbx1* expression together with a set of genes which are sequentially upregulated during olfactory neurogenesis. Thus, early progenitors committed to a neuronal fate express the proneural gene *Ascl1* (formerly known as *Mash1*), they generate immediate neuronal precursors (INPs) identified by *Neurog1* expression, which undergo terminal differentiation in postmitotic neurons expressing *Stmn2* (*Scg10*), *Gap43* and finally *Ncam1* according to their maturation state (Calof et al., 2002). Double fluorescent *in situ* hybridization experiments performed at E11.5 indicated that *Dbx1* mRNA never colocalized with either *Ascl1, Stmn2, Gap43* or *Ncam1* (Fig 1D). By contrast, we found all *Dbx1*-expressing cells to be *Neurog1*^+^ (Fig 1D). We therefore concluded that *Dbx1* expression is temporally regulated during the process of olfactory neurogenesis, restricted to a subset of INPs. We then took advantage of the fact that in *Dbx1*^*LacZ/+*^ embryos, βGal persists longer than the Dbx1 protein, resulting in short term tracing (Pierani et al., 2001). Immunostaining on these embryos indicated that β-Gal colocalizes with Lhx2, which is expressed by INPs and maintained in neurons, HuC/D, an early neuronal marker, and CTIP2, an early marker of OSN, but is almost never found in Sox2^+^ progenitor cells (Fig 1E). Taken together, these results indicate that *Dbx1* is expressed by INPs just prior to neuronal differentiation (Fig 1F).

A strong Dbx1 expression was also found in adults as illustrated by X-Gal staining of *Dbx1*^*LacZ/+*^ animals (Fig 2A,B). The pattern of staining, with sharp boundaries between positive and negative regions in the OE, indicated that Dbx1 expression is zonally restricted at this stage (Fig 2B). In order to identify the cells that express Dbx1, we stained the olfactory epithelium with antibodies against Sox2 and CTIP2 to distinguish between glial sustentacular (SUS) cells (Sox2^+^) on the apical side and the neuronal compartment (CTIP2^+^) on the basal side (Fig 2C). We found that 71% ± 6% βGal^+^ cells are neurons and the remaining are SUS cells (Fig 2E). In addition, immunostaining revealed that approximately half of βGal^+^ cells (46% ± 5%) are mature olfactory sensory neurons (OSNs) expressing the olfactory marker protein (OMP) (Fig 2D, E). We therefore concluded that *Dbx1* is expressed during both embryonic and adult olfactory neurogenesis as well as during adult gliogenesis.

**Figure 2.**
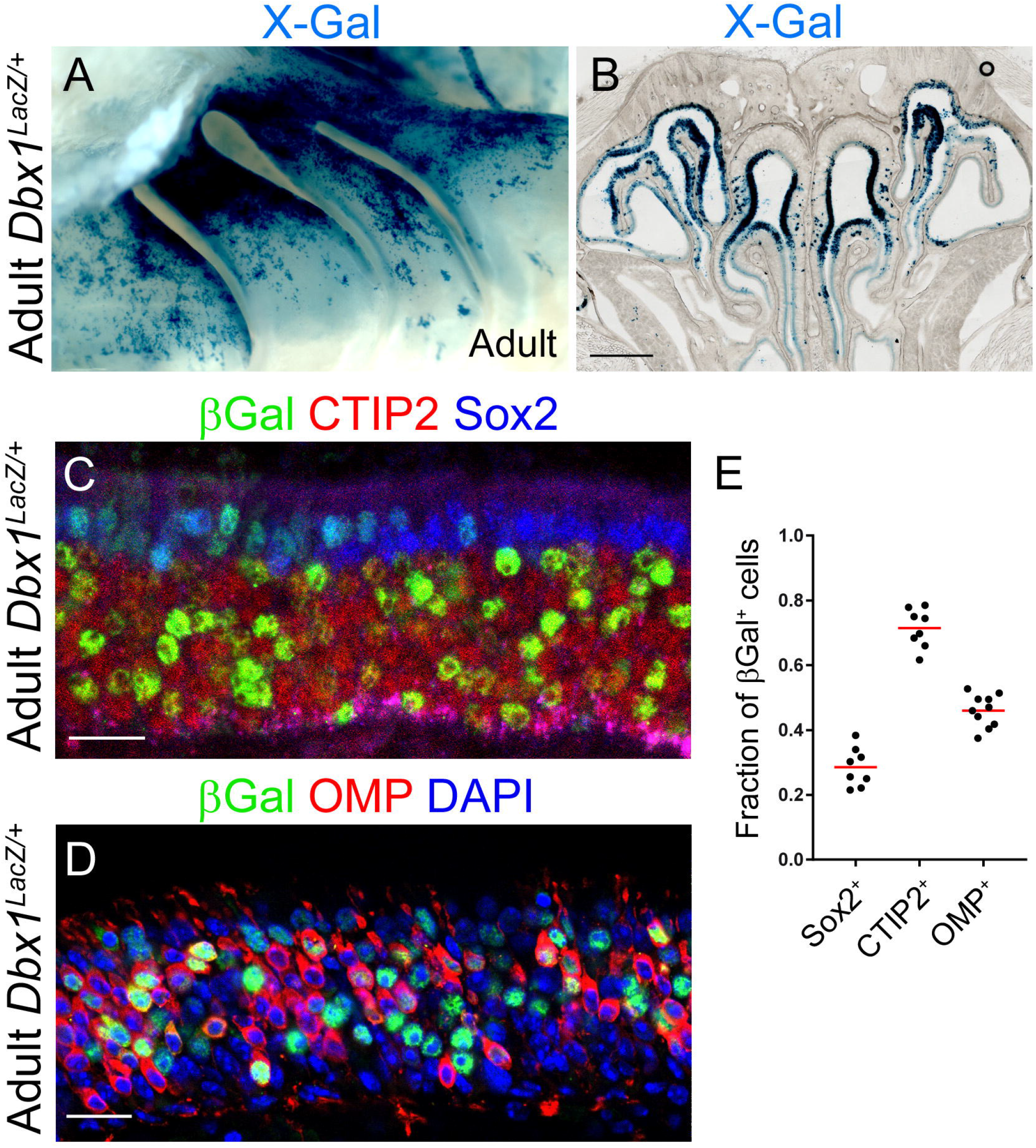
Dbx1 expression in the adult olfactory epithelium. (A) Whole-mount X-Gal staining of the olfactory epithelium of an adult *Dbx1*^*LacZ/+*^ mouse. (B) X-Gal staining on a coronal section through the olfactory epithelium of an adult *Dbx1*^*LacZ/+*^. (C) Immunostaining on the olfactory epithelium of an adult *Dbx1*^*LacZ/+*^ mouse using antibodies against βGalactosidase (green), CTIP2 (red) and Sox2 (blue). (D) Immunostaining on the olfactory epithelium of an adult *Dbx1*^*LacZ/+*^ mouse using antibodies against βGalactosidase (green) and OMP (red), as well as DAPI to stain nuclei (blue). (E) Quantification of the percentage of marker-positive cells among βGalactosidase-positive cells. Scale bars: 1mm in B, 20μm in C, D.

### Dbx1 is dispensable for olfactory neurogenesis

In order to investigate the consequences of *Dbx1* loss on olfactory neurogenesis, we analyzed *Dbx1*^*LacZ/LacZ*^ animals and compared them to *Dbx1*^*LacZ/+*^ counterparts. Because mutant animals die at birth (Pierani et al., 2001), analyses could only be conducted up to late embryonic stages. In mutants, not only the intensity of βGal staining but also the number of βGal^+^ cells in the developing OE is increased compared to heterozygous littermates (Fig 3A, C, D and data not shown). This could result from (i) increased proliferation of βGal^+^ cells, (ii) decreased cell death or (iii) increased βGal accumulation due to the presence of a second LacZ allele. We first performed PH3 immunostaining to label mitotic cells, and found no significant differences between control and mutants, either in the distribution or number of positive cells (57 ± 9 per mm of OE and 54 ± 13 respectively; Fig 3A-A’’). To detect apoptosis, we performed immunostaining against activated Caspase3. In both mutant and heterozygous animals, we found very few apoptotic cells (9 ± 3 per mm of OE and 7 ± 4 respectively) indicating that the loss of *Dbx1* does not impair cell viability (Fig 3B-B’’). We then tested whether the loss of *Dbx1* could impact cell identity. Similar to heterozygous controls, we found that mutant βGal^+^ cells were Sox2^-^, Lhx2^+^ and CTIP2^+^ (Fig 3C), indicating that they retain their identity of neurogenic progenitor despite the loss of *Dbx1*. We further analyzed olfactory neurogenesis in mutant and control animals. Tuj1 staining appeared similar in control and mutant OE, including in the area of βGal^+^ cells (Fig 3D, D’). The number and trajectory of migrating GnRH^+^ neurons was also unchanged at E12.5 (Fig 3E, E’). Bundles of sensory axons growing from the OE to the brain, identified by Peripherin or Tuj1 staining, appeared normal in thickness and position in *Dbx1*^*LacZ/LacZ*^ animals (Fig 3F, F’) suggesting that OSN/VSN axon guidance does not require Dbx1. This was confirmed by OMP staining at E14.5, which allowed to visualize cell bodies of mature OSN (Fig 3G, G’) and VSN (Fig 3I, I’) in the OE and VNO, respectively, as well their axons reaching the olfactory bulb (Fig 3H-H’). No qualitative or quantitative difference (Fig 3G’’) was found between control and mutants.

**Figure 3.**
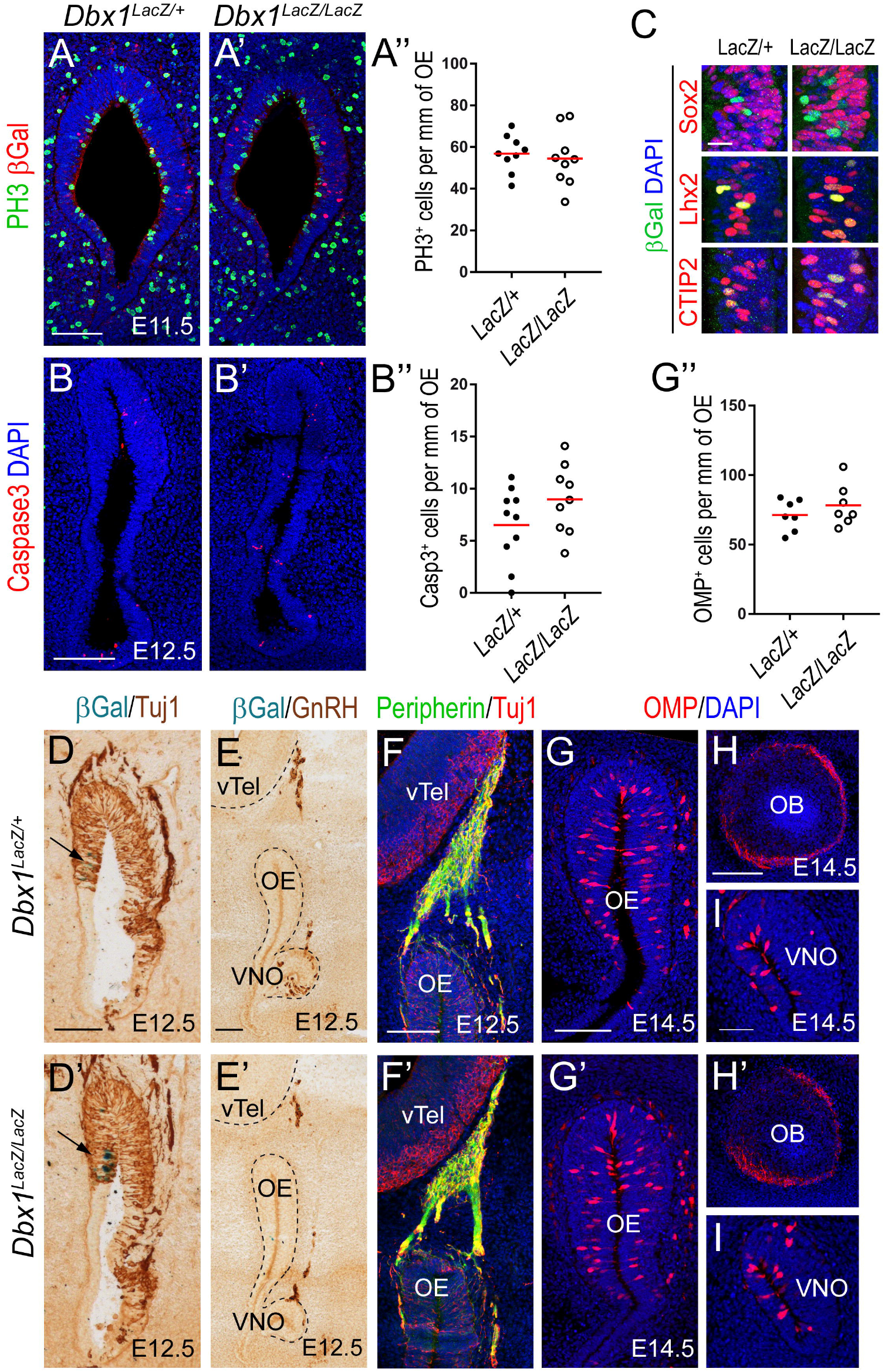
Consequences of Dbx1 loss during olfactory neurogenesis. (A, A’) Immunostaining against phosphor-histone 3 (green) and βGalactosidase (red) on coronal cryosections of the olfactory pit in control *Dbx1*^*LacZ/+*^ (A) and mutant *Dbx1*^*LacZ/LacZ*^ (A’) E11.5 embryos. (A’’) Quantification of mitotic PH3^+^ cells. (B, B’) Immunofluorescence against activated caspase 3 (red) in control *Dbx1*^*LacZ/+*^ (B) and mutant *Dbx1*^*LacZ/LacZ*^ (B’) E11.5 embryos. (B’’) Quantification of apoptotic Caspase 3^+^ cells. (C) Immunostaining for βGalactosidase (green) and either Sox2, Lhx2 or CTIP2 (red) in *Dbx1*^*LacZ/+*^ and *Dbx1*^*LacZ/LacZ*^ embryos. (D) DAB immunostaining against Tuj1 (brown) combined with X-Gal staining (cyan) in *Dbx1*^*LacZ/+*^ (D) and *Dbx1*^*LacZ/LacZ*^ (D’) E12.5 embryos. (E) Same as D using GnRH immunostaining. (F) Immunofluorescence against Peripherin (green) and Tuj1 (red) in control (F) and mutant (F’) embryos. (G-I) OMP immunostaining showing the E14.5 OE (G), OB (H), and VNO (I). (G’’) Quantification of OMP^+^ cells. vTel: ventral telencephalon, OE: olfactory epithelium, VNO: vomeronasal organ, OB: olfactory bulb. Scale bars: 100μ in A, B, D, E, F and G, 20μm in C, 200μ in H, 50μm in I.

In order to assess terminal neuronal differentiation, we monitored the expression of vomeronasal and olfactory receptor genes in control and mutant animals at E18.5 (Fig 4A). In the VNO, neurons expressing receptors from the V1R family (Dulac and Axel, 1995) occupy apical positions whereas those expressing V2R genes (Herrada and Dulac, 1997; Matsunami and Buck, 1997; Ryba and Tirindelli, 1997) are located more basally. *In situ* hybridization experiments using probes targeting multiple genes of either family indicated that such an organization is maintained in Dbx1 mutants (Fig 4B,C). In the OE, the expression of olfactory receptor genes is subjected to zonal restriction such that OSN expressing the same receptor are confined to the same region (Ressler et al., 1993; Vassar et al., 1993; Zapiec and Mombaerts, 2020). At E18.5, *Omp* expression indicates that the number and distribution of mature OSN are comparable in mutants and control littermates (Fig 4D, D’), consistent with our findings at E14.5. In addition, the zonal organization of the OE appears unaffected by *Dbx1* loss as illustrated by the consecutive expression of *Olfr247, Olfr1395, Olfr449* and *Olfr1507* in consecutive dorsomedial to ventrolateral domains (Fig 5E-H).

**Figure 4.**
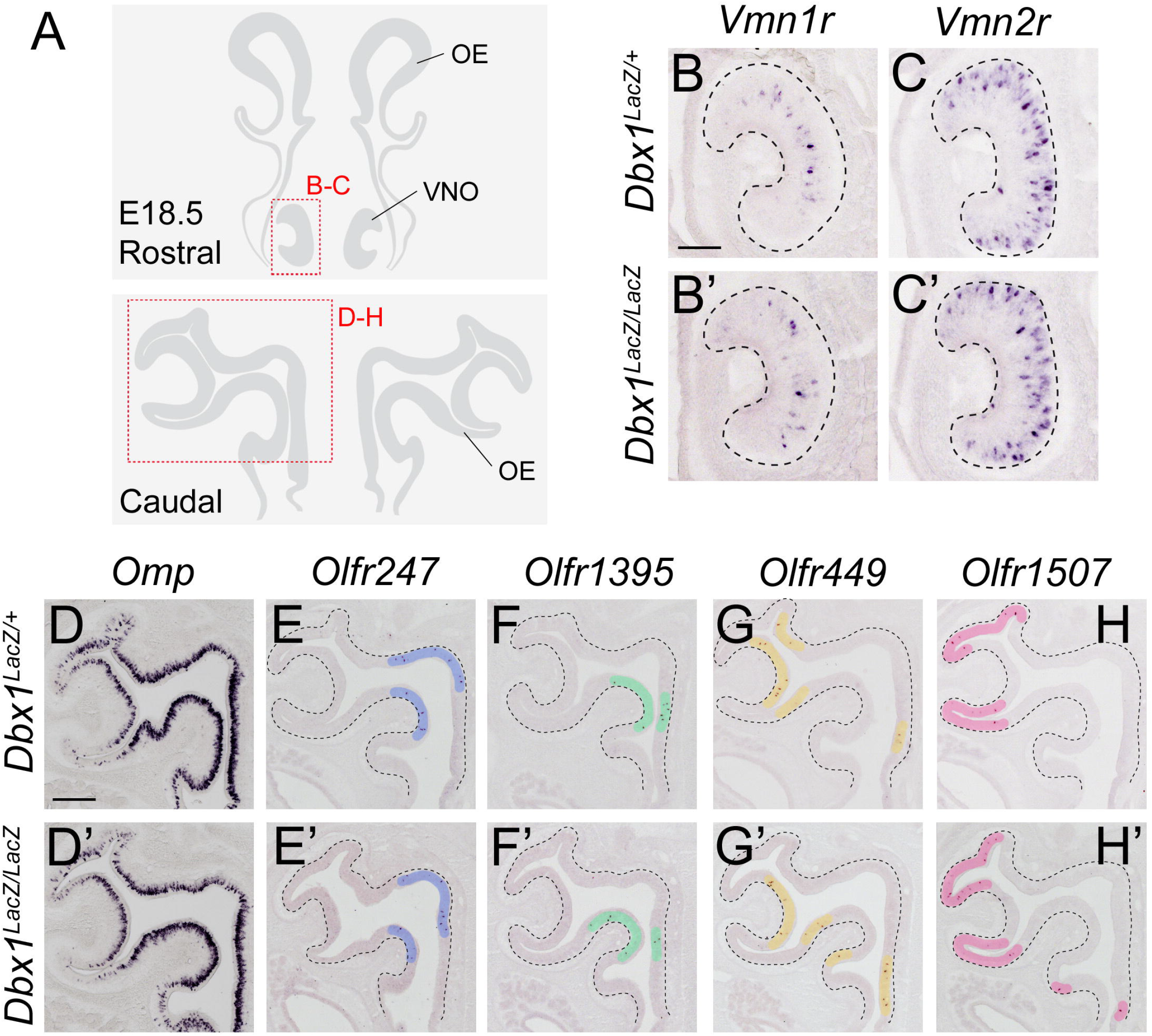
Consequences of Dbx1 loss on the zonal organization of the olfactory system. (A) Drawing of the position corresponding to the distinct panels of the figure. (B) *In situ* hybridization using a probe against *Vmn1r* genes in *Dbx1*^*LacZ/+*^ (B) and *Dbx1*^*LacZ/LacZ*^ (B’) embryos. (C) Same as (B) using a probe against *Vmn2r* genes. (D-H) *In situ* hybridization using probes against *Omp* (D), *Olfr247* (E), *Olfr1395* (F), *Olfr449* (G) and *Olfr1507* (H) in *Dbx1*^*LacZ/+*^ (D-H) and *Dbx1*^*LacZ/LacZ*^ (D’-H’). OE: olfactory epithelium, VNO: vomeronasal organ. Scale bars: 100μm in B, 200μm in D.

**Figure 5.**
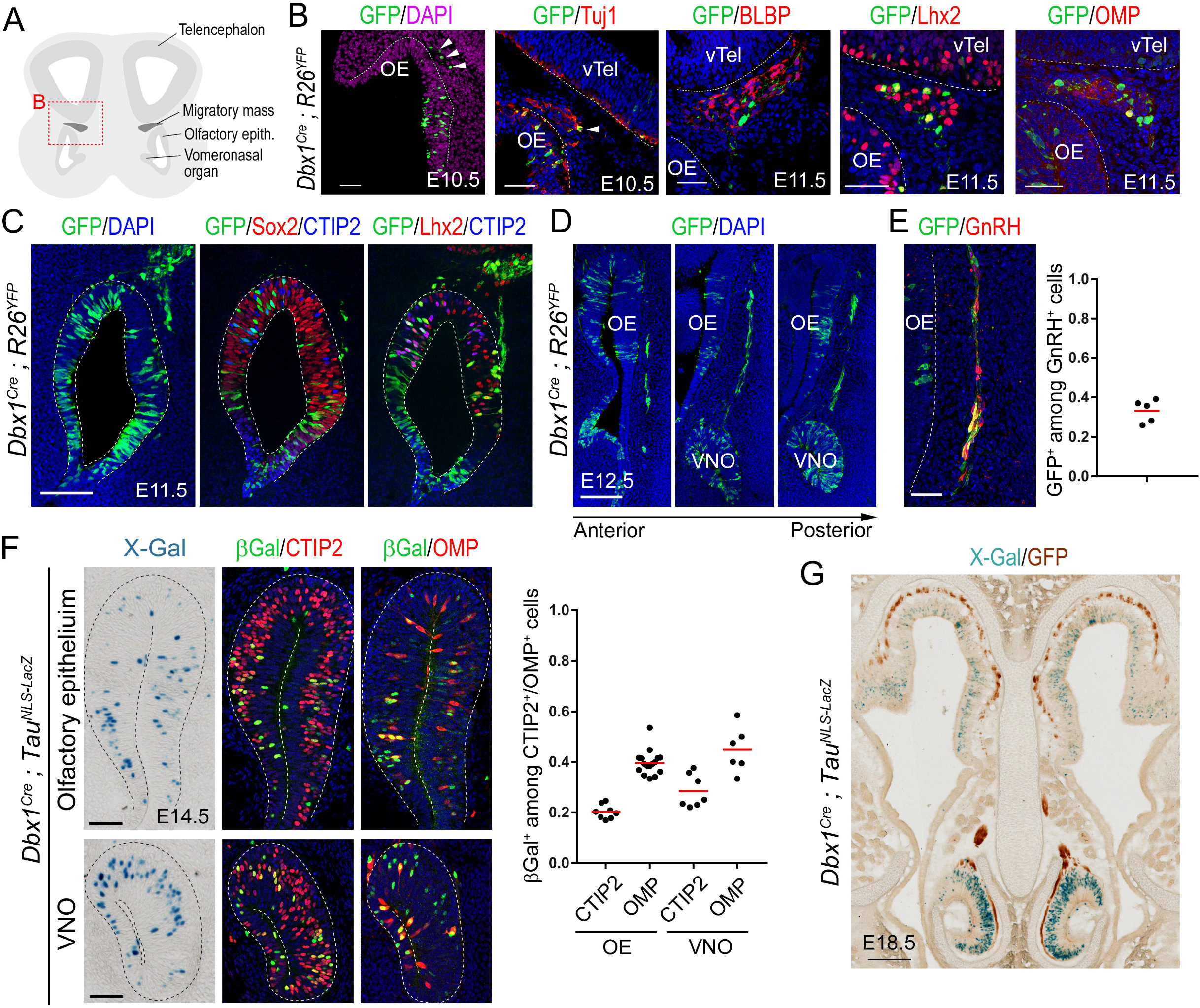
Genetic tracing of the Dbx1 lineage in olfactory epithelium derivatives. (A) Drawing of the region of the migratory mass shown in (B). (B) Immunofluorescence against GFP (green) and either Tuj1, BLBP, Lhx2 or OMP (red) in *Dbx1*^*Cre*^*;Rosa26*^*YFP*^ embryos of the indicated stage. DAPI is shown in blue or purple. (C) Immunofluorescence for GFP (green), Sox2 or Lhx2 (red) and DAPI or CTIP2 (blue) at E11.5. (D) Immunostaining of GFP along the antero-posterior axis of the olfactory system at E12.5. (E) Co-immunostaining for GFP (green) and GnRH (red), and quantification of the overlap. (F) X-Gal staining (cyan) and immunofluorescence for βGalactosidase (green) and CTIP2 or OMP (red) in the OE (top) and VNO (bottom) of E14.5 *Dbx1*^*Cre*^*;Tau*^*YFPNLS-LacZ*^ embryos. Quantification of the colocalisation is shown. (G) X-Gal staining (cyan) combined with DAB staining for GFP (brown) at E18.5 in *Dbx1*^*Cre*^*;Tau*^*YFPNLS-LacZ*^ embryos. vTel: ventral telencephalon, OE: olfactory epithelium, VNO: vomeronasal organ. Scalle bars: 50μ in B, E and F, 100μm in C and D, 200μm in G.

Taken together our data suggest that the production and maturation of the various neuronal subtypes in the developing olfactory system is not dramatically affected in the absence of *Dbx1*.

### Dbx1 lineages contribute to multiple components of the olfactory system

Given *Dbx1* expression in olfactory neural progenitors, we decided to explore the contribution of the Dbx1 lineage to the various components of the olfactory system. We performed genetic tracing using *Dbx1*^*Cre*^*;R26*^*YFP*^ animals (Griveau et al., 2010) in order to permanently label the progeny of Dbx1-expressing progenitor cells. At E10.5, consistent with the early onset of D*bx1* expression, scattered GFP^+^ cells were found in the developing OE (Fig 5A,B). In addition, GFP^+^ cells were detected in the mesenchyme between the OE and the ventral telencephalon, an area containing a group of cells referred to as the migratory mass (MM) (Miller et al., 2010; Valverde et al., 1992). The MM was previously described as a heterogeneous population, containing both neuronal and glial cells, and proposed to act as a guidepost for sensory axons growing towards the brain (Miller et al., 2010). We found that the Dbx1 lineage exclusively contributes to the neuronal content of the developing MM as GFP^+^ cells are also positive for Tuj1, which labels neurons, but never for BLBP, a marker for the glial olfactory ensheathing cells (Fig 5B). Consistently, most GFP^+^ cells in the MM were found to be Lhx2^+^ whereas a fraction expressed OMP (Fig 5B). At E11.5 GFP^+^ cells were found in both the OE and the respiratory epithelium (Fig 5C). Consistent with our Dbx1 expression data, most Dbx1-derived cells in the OE were found to express Lhx2 and CTIP2 but rarely Sox2, indicative of their neuronal (rather than progenitor) identity. At E12.5, Dbx1-derived cells were found in the vomeronasal organ (VNO) (Fig 5D). In addition, GFP^+^ cells were observed in the mesenchyme medial to the OE, forming streams of cells displaying a morphology highly reminiscent of neurons migrating from the VNO towards the brain. Most but not all of these cells could be identified as GnRH^+^ neurons (Fig 5E). We found that the Dbx1-lineage accounts for approximatively one third of all GnRH neurons (33% ± 6%).

From E14.5 onwards, it is possible to genetically label specifically the neuronal progeny of Dbx1-expressing cells using *Dbx1*^*Cre*^*;Tau*^*NLS-LacZ*^ animals. Dbx1-derived neurons are evident in both the OE and VNO at this stage (Fig 5F). Quantifications indicated that in the OE, the Dbx1 lineage accounts for 20% ± 3% of immature CTIP2^+^ neurons and 40% ± 5% of fully differentiated OMP^+^ OSNs, suggesting a decrease with time in the contribution of the Dbx1 lineage to olfactory neurogenesis. Similarly, in the VNO, 28% ± 7% of immature neurons and 45% ± 9% of mature vomeronasal sensory neurons (VSNs) derive from Dbx1-expressing progenitors (Fig 5F). At E18.5 the distribution of Dbx1-derived neurons within the OE and VNO was found to be homogenous (Fig 5G).

Taken together, these data indicate that Dbx1-expressing progenitors contribute significantly to the various neuronal components of the peripheral olfactory system, and support the idea that OSNs, VSNs and GnRH neurons are heterogeneous populations.

### Dbx1^+^ progenitors remain neurogenic in Mash1 mutants

Previous studies concur to the idea that the proneural transcription factor Ascl1 is a key player in olfactory neurogenesis (Sokpor et al., 2018). In the OE of *Ascl1* mutants, the expression of *Ngn1* is almost completely lost and that of *Lhx2* severely reduced (Cau et al., 1997; Cau et al., 2002). This results in a dramatic reduction of OMP^+^ OSN at later stages (Guillemot et al., 1993). To date, the identity of *Ascl1*-independent OSN remain elusive. We therefore thought to analyze *Dbx1* expression in *Ascl1* mutants. Surprisingly, the number and distribution of *Dbx1*^+^ cells appeared unaffected by the loss of *Ascl1* (Fig. 6A). Furthermore, double *in situ* hybridization experiments indicated that *Dbx1*^+^ cells remain *Ngn1*^+^ and *Lhx2*^+^ in *Ascl1*^-/-^ embryos (Fig. 6B,C), indicating they retain an identity of neurogenic progenitor. Immunostaining confirmed this result at the protein level, and further indicated that Lhx2^+^ cells remaining in *Ascl1* mutants are located nearby Dbx1-expressing regions (Fig. 6D). Therefore, opposite to the rest of the developing OE, the expression of *Ngn1* in *Dbx1*^+^ cells is independent of *Ascl1*, and Dbx1-expressing progenitors appear able to undergo Ascl1-independent neurogenesis. Finally, since Dbx1-expressing progenitors also contribute to the GnRH lineage, we investigated GnRH neuron production in *Ascl1* mutants and found it is severely reduced, but not completely abolished, confirming previously published data (Taroc et al., 2020; Tucker et al., 2010) and reminiscent of the ∼1/3 contribution of the Dbx1 lineage to GnRH neurons. This strongly suggests that Dbx1^+^ progenitors account for the *Ascl1*-independent specification of GnRH neurons and further underlines the extent of diversity among progenitors involved in olfactory neurogenesis.

**Figure 6.**
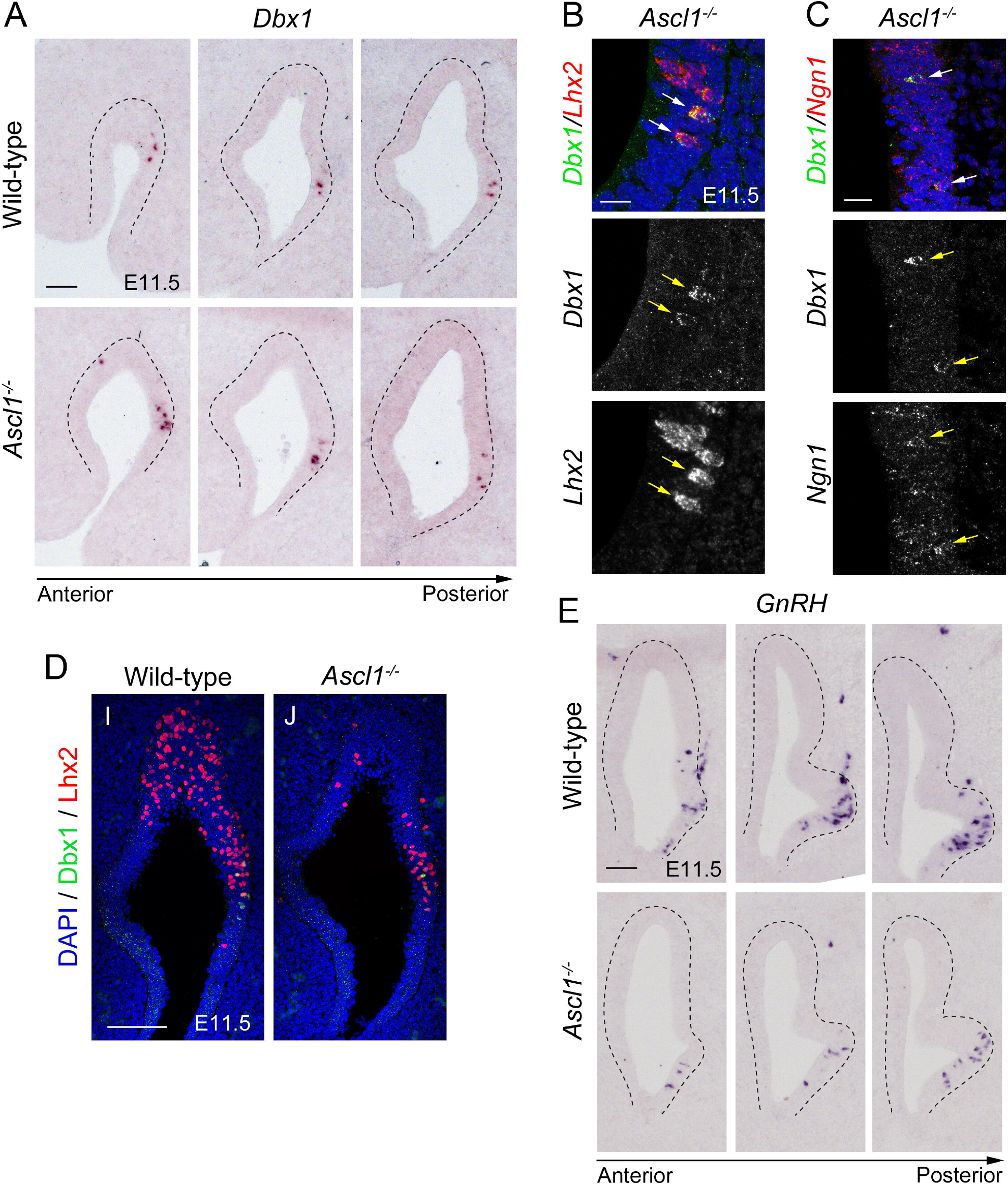
Consequences of Ascl1 loss for Dbx1-related olfactory neurogenesis. (A) *In situ* hybridization for *Dbx1* along the antero-posterior axis of the E11.5 olfactory epithelium in control and *Ascl1*^*-/-*^ embryos. (B) Double fluorescent *in situ* hybridization for *Dbx1* (green) and *Lhx2* (red) in the olfactory epithelium of E11.5 *Ascl1*^*-/-*^ embryos. (C) Double fluorescent *in situ* hybridization for *Dbx1* (green) and *Neurog1* (red) in E11.5 *Ascl1*^*-/-*^ embryos. (D) Immunostaining for Dbx1 (green) and Lhx2 (red) in control and *Ascl1*^*-/-*^ E11.5 embryos. (E) In situ hybridization for *GnRH* along the antero-posterior axis of the E11.5 olfactory epithelium in control and *Ascl1*^*-/-*^ embryos. Scale bars: 50μm in A and E, 20μm in B and C, 100μm in D.

## Discussion

### Dbx1 in the sensory olfactory epithelium

In the spinal cord, Dbx1 acts as a fate selector, expressed in progenitors and imposing V0 interneuron identity : in its absence, neurons fail to acquire their normal identity and adopt the alternative V1 fate (Pierani et al., 2001). By contrast, our data suggest that in the olfactory sensory system, Dbx1 does not act as a strong fate inducer. Although we cannot formally exclude the possibility that neurons derived from *Dbx1*^*LacZ/LacZ*^ progenitors display some degree of misspecification, we found that expression of cardinal markers, global organization of the OE/VNO, axon targeting of OSN/VSN and GnRH neuron migration appeared normal. The only noticeable difference concerned the increased number of βGal^+^ cells observed in knock-outs. One possible explanation is that two LacZ alleles allow increased accumulation of βGal. An interesting alternative hypothesis is that Dbx1 is involved in the cycling behavior of progenitors. Consistently, Dbx1 gain-of-function was proposed to induce terminal differentiation (García-Moreno et al., 2018).

One interesting feature of Dbx1 expression in the spinal cord and forebrain is its typical position at borders between domains (Bielle et al., 2005; Lu et al., 1992; Mangale et al., 2008). Tissue patterning within the OE was shown to occur, consisting in the graded expression of transcription factors, and perturbation of such gradients was shown to affect progenitor proliferation properties and precursor identities (Tucker et al., 2010). Our data therefore support a model in which progenitors of the olfactory epithelium form a continuum from which distinct classes of neurons emerge, similar to what we have recently described along the dorso-ventral axis of the mouse pallium (Moreau et al., 2021). Accordingly, Dbx1 expression would be constrained at specific coordinates within this gradient, corresponding to transition zones, and the corresponding progeny participate in multiple lineages, thus revealing an otherwise hidden degree of progenitor diversity. Whether such progenitor diversity translates into distinct neuronal identities awaits further studies.

### Diversity among sensory olfactory derivatives

Distinct levels of heterogeneity within OSN, VNO and GnRH have already been documented. One of the best example relates to the zonal expression of odorant receptors (Zapiec and Mombaerts, 2020). Currently, the molecular rationale allowing to confine the expression of odorant receptor genes to specific zones of the OE remains unclear. Adult expression of Dbx1 appear to follow such a zonal organisation, with relatively sharp boundaries between regions of high and low/absent expression. Lineage tracing using Nestin-Cre mice was also shown to result in zonal pattern of recombination (Murdoch and Roskams, 2008), and multiple homeodomain transcription factors are expressed in a zonally restricted manner (Parrilla et al., 2016). This point to the idea that the zonal organization of the OE could reflect OSN ontogeny.

Diversity among GnRH neurons has also been demonstrated at the morphological level (Cottrell et al., 2006; Wray and Hoffman, 1986), on molecular bases (Constantin et al., 2009; Jasoni et al., 2005; Klenke et al., 2010), or related to the birthdate and final position of the cells (Jasoni et al., 2009). Furthermore, lineage tracing previously argued in favor of multiple embryonic origin for GnRH neurons, as approximatively one third was shown to derive from a neural crest lineage and the remaining two thirds from ectodermal lineage (Forni et al., 2011). Whether Dbx1-derived and neural crest lineages overlap or correspond to distinct GnRH neuron subtypes remains to be established. Dbx1 mRNA and protein expression argues in favor of the local specification of Dbx1 lineage in the OE. Nevertheless, the previous description we made of early (E8.5) Dbx1-expressing progenitors contributing to a subset of cranial neural crests cells (Causeret et al., 2011) could support a certain degree of overlap between the two lineages.

Furthermore, a recent preprint investigating the cellular diversity and connectivity patterns in the accessory olfactory system reported that Dbx1-driven diversity is not only observed in the VNO but also in the accessory olfactory bulb and medial amygdala (Prakash et al., 2022). This work points to the idea of a molecular code involved in the assembly of olfactory circuits that would be conserved from the sensory epithelia to the central processing structures.

### *Ascl1*-independent neurogenesis in the olfactory epithelium

Ascl1 is a proneural transcription factor and, as such, promotes neuronal differentiation in the olfactory epithelium (Cau et al., 1997; Cau et al., 2002; Guillemot et al., 1993; Murray et al., 2003). Upon *Ascl1* loss, OSN, VSN and GnRH differentiation is severely affected, with only few neurons reaching a mature state (Guillemot et al., 1993; Murray et al., 2003; Taroc et al., 2020). The origin of these few neurons so far remained elusive. We show that Dbx1-expressing cells in the olfactory epithelium of *Ascl1* mutants retain a neurogenic potential, as they maintain *Ngn1* expression. We therefore postulate that neurons produced independently of *Ascl1* belong to the Dbx1 lineage. This hypothesis is supported by the observation that the remaining OMP^+^ cells in *Ascl1* mutants are early born (Tucker et al., 2010). In addition, the same authors calculated that 29% of Tuj1^+^ neurons are maintained in the E11.5 mutant OE, matching the contribution of the Dbx1 lineage we describe.

One remaining open question is the identity of Dbx1^+^ precursors. They could either derive from *Ascl1*-expressing progenitors, in which case one could postulate that *Dbx1* is able to compensate for the loss of *Ascl1*. An alternative possibility is that *Dbx1*^+^ progenitors represent a previously uncharacterized progenitor pool, distinct from the *Ascl1* lineage, and therefore insensitive to *Ascl1* loss.

Overall, our work points to the diversity that exists in the olfactory sensory system, and raises new questions regarding how progenitor and neuron diversity are generated.

## Acknowledgements

The authors wish to thank the animal facility of the Institut Jacques Monod. We acknowledge the ImagoSeine facility, member of the France BioImaging infrastructure supported by the French National Research Agency (ANR-10-INSB-04, “Investments for the future”) for help with confocal microscopy. We are grateful to Vanessa Ribes for the kind gift of antibodies, Adrien Calvairoly for help with collecting *Ascl1* mutants and members of the Pierani laboratory for helpful discussions. FC is an Inserm researcher. AP is a CNRS investigator. This work was supported by grants from Agence Nationale de la Recherche (ANR-2011-BSV4-023-01), FRM (Fondation pour la Recherche Médicale) (INE20060306503 and «Equipe FRM DEQ 20130326521») to AP.

## Authors contribution

Conceptualization: FC

Methodology: FC, AC, AP

Investigation: FC, MF, MXM, EN, RO, AC

Resources: CP

Writing – original draft preparation: FC

Writing – review and editing: FC, RO and AC

Visualization: FC, MF, MXM, EN

Supervision: FC

Project administration: FC

Funding acquisition: AP

## Competing interests

No competing interests declared.

## Data and materials availability

All data are available in the main text or the supplementary materials.

## Notes

### Competing Interest Statement

The authors have declared no competing interest.

